# Genetic variation of the effects of spontaneous mutation on size at birth

**DOI:** 10.64898/2026.01.29.702524

**Authors:** Matthew R. Bruner, Trenton C. Agrelius, Krista B. Harmon, Jeffry L. Dudycha

## Abstract

Spontaneous mutation underlies all genetic variation, and thus influences the evolutionary dynamics of complex traits. Although much work has estimated mutation rates for fitness or at molecular scales, we have comparatively little information about mutational influences on other complex phenotypes. We conducted four mutation accumulation experiments with independent clones of *Daphnia pulex*, and then measured the effect of spontaneous mutation on size at birth, a complex trait whose connection to fitness depends on ecologically-mediated tradeoffs. Therefore it is unclear whether mutations, which are usually neutral or deleterious, should decrease or increase size at birth. In two experiments, individual instances of increased size at birth were common, whereas in the other two experiments, instances of decreased size at birth were common. Together, our data show that genetic background is an important determinant of the consequences of mutation for complex traits, and that mutation rates and direction can vary within species.

## Introduction

Genetic variation within and among populations originates through spontaneous mutation. For complex traits influenced by many loci, we can gain a better understanding of how they evolve if we know the effects of mutation on the mean and variance of the trait, even in the absence of knowing the specific genes involved. Mutation accumulation (MA) experiments approach this problem via experimental evolution (Lynch, et al. 1999; Halligan & Keightley 2009). In these experiments, replicate lines are derived from an individual ancestor (or male-female pair of ancestors), and propagated through single-individual (or -pair) descent. In this way, effective population size is minimized, lines are shielded from most but not all selection, and spontaneous mutations accumulate generation after generation. After some period of time, genetic- *i.e.*, mutational- variation of a trait can be measured among replicate lines. If an unevolved ancestor is available, mutational means can be compared between the ancestor and MA lines to establish directionality.

Size at birth is a critical life history trait that establishes the trajectory for an individual’s future fitness (Stearns 1992; Roff 2002). It predicts subsequent life history traits and physiology in both vertebrates and invertebrates (Ebert 1991; Oksanen, et al. 2003, 2007; Mappes & Koskela 2004; Ylonen, et al. 2004; Bashey 2008; Schrader & Travis, 2012; King, et al. 2016). The general concept is that if an individual is born small, it has few resources to draw on at the start of juvenile development and often has limits on its ability to acquire resources. More resources may be required for the individual to reach reproductive maturity, at the cost of delayed maturation or possibly reduced size at maturation, which may in turn limit reproductive fitness (Stearns 1992; Roff 2002). Individuals born large have contrasting advantages in resource acquisition and reaching maturity. While smaller individuals can recover some mass through “catch-up growth,” rapid accumulation of mass can have detrimental effects on fitness (Arendt 1997; Singhal 2017). Usually, larger size at birth is advantageous to the individual, but not necessarily to the mother that produced the individual, who faces a trade-off between offspring size and number (Smith & Fretwell 1974; Stearns 1992; Roff 2002).

We sought to quantify the effects of mutation on size at birth with MA experiments in the crustacean *Daphnia pulex*, an ideal model system that can be propagated asexually and in which the consequences of variation of size at birth are understood well (Ebert 1991; Tessier & Consolatti 1991; Tessier, et al 1992; Lampert 1993; Yampolsky & Scheiner 1996; Sawinska 2002, 2004; Reede, 2003; Dudycha & Lynch 2005; Dudycha, et al. 2013). Since the effects of mutation on complex traits can depend on the complement of alleles an individual already carries, we conducted experiments starting with four independent ancestors.

## Materials and Methods

### Study system

*Daphnia pulex* are freshwater microcrustaceans often found inhabiting small, temporary ponds. Their typical reproductive system is cyclical parthenogenesis, and under laboratory conditions they can be induced to reproduce asexually indefinitely, allowing a large number of genetically identical individuals to be generated. Every few days, adult females release a clutch of eggs into their brood chamber, which then develops and is released as neonates when the adult molts at the end of the instar. With typical *Daphnia*, haploid eggs can be produced in response to environmental cues, and following fertilization, these are released in a structure called an ephippium where they remain dormant until suitable hatching cues are perceived.

### Mutation accumulation lines

In May, 2016 we launched four identical mutation accumulation experiments with obligately asexual clones of *Daphnia pulex*. Unlike those with the typical life cycle, in these clones meiosis is always suppressed in females and therefore dormant eggs are produced asexually (Hebert 1981; Hebert & Finston 2001; Eads, et al. 2012; Ye, et al. 2021). Each experiment began with a distinct clone isolated from a separate population. The four clones are LIS3, TRO3, MORG5, and SED2, all of which were confirmed obligately asexual (Paland & Lynch 2006; for SED2: [Authors], unpubl. data) and have had their genomes sequenced (Tucker et al 2013; Li, et al 2014; GenBank Accessions: MORG5, SAMN02252737; SED2, SAMN02252738; TRO3, SAMN02252729; LIS3, SAMN02252731). The clones were originally isolated from ponds widely spread across the North American range of *D. pulex*, from Maine (TRO3: 44° 38’ N, 69° 14’ W), Ontario (LIS3: 43° 44’ N, 80° 57’ W), Michigan (MORG5: 42° 13’ N, 83° 44’ W), and Minnesota (SED2: 46° 56’ N, 92° 13’ W). Although we have not specifically assessed genetic variation among our ancestral clones, *Daphnia pulex* populations are generally observed to be strongly divergent (Crease, et al. 1990; Pfrender, et al. 2000; Straughan & Lehman 2000; Maurki, et al 2022) and phylogeographic studies of obligate asexuality suggest that the geographic scales separating our populations reflect substantial genetic divergence (Paland, et al 2005; Crease, et al. 2012; Tucker, et al. 2013, Xu, et al. 2015).

To start the experiments, we isolated thirty female neonates from a single adult female and placed each neonate into separate 150-ml beakers filled with 100 ml filtered lakewater. Each isolated neonate became the first generation of a replicate MA line. We propagated lines by haphazardly choosing a single offspring of the current generation, transferring it to a new beaker, and labeling it as the next generation. Normally, offspring were nearing maturity at the time of a transfer. To avoid biasing transfer against individuals least able to escape the experimenter’s pipette, the person performing transfers captured a targeted individual. To avoid biasing transfers for the most active or largest individuals, experimenters usually aimed to transfer the second individual that was noticed when first observing a beaker. We advanced generations approximately every two weeks. At each transfer two “sisters” of the focal individual were separately reserved in a 50-ml centrifuge tube to serve as back-ups in case the focal individual was male, or never produced female offspring. As an additional backup, the two prior generations were kept at 10°C. In some instances, we restored lines that had gone extinct by hatching asexually produced dormant eggs collected from a prior generation. The number of generations that have passed separates over the course of the experiment due to the cumulative effects of producing male or dormant offspring. This system guards against accidents while minimizing effective population size and limiting most effects of selection. However, mutations with large deleterious effects when heterozygous (e.g., dominant lethals) cannot be captured in this type of experiment.

We propagated our MA lines under standard lab conditions for rearing *Daphnia*: filtered lakewater (to 1μm), 20°C, 12:12 light:dark photoperiod, fed vitamin-fortified *Ankistrodesmus falcatus* daily, and surface tension reduced with cetyl alcohol. For additional details see Davenport, et al. (2021).

### Ancestral controls

Each experiment is comprised of the set of mutation accumulation lines, plus asexually produced ephippia of the ancestor that were induced at the start of the experiment and preserved at −80°C. The ancestors’ eggs serve as unmutated controls in phenotypic assays. We used additional daughters of each individual ancestor to grow a large population and then induced dormant egg production via crowding. This way, we collected >1000 dormant eggs that we estimate are three to four generations removed from the individual stem mother of each experiment. Immediately prior to a phenotypic assay, we hatched ephippia by gradually exposing them to increased temperature and light (details in Davenport, et al. 2021). Hatching success rates were typically ∼75%. Our aim was to hatch enough ephippia for replicate control lines to be in at least a 1:2 ratio with MA lines.

### Size at birth assays

The four sets of MA lines were assayed sequentially rather than concurrently due to operational constraints. Because of this and differences among sets in their reproductive characteristics, the assays were performed at different generations for the different sets of lines. Information on the timing and replication in assays of each set of MA lines can be found in Table 1. Some MA lines were lost prior to our size at birth assays. These losses primarily occurred when all living females of a line (focal individual + backups) produced only dormant or male offspring. The SED lines, which were most prone to producing male and dormant offspring (Table 1), suffered notably more losses than the other sets.

**Table 1.** Comparative information for four independent mutation accumulation experiments.

|  | MORG | SED | TRO | LIS |
| --- | --- | --- | --- | --- |
| Assay start date | 16 DEC 20 | 07 JUL 20 | 15 OCT 20 | 06 AUG 19 |
| Mean # of generations | 93.4 | 72.3 | 86.5 | 52.2 |
| # of MA lines assayed | 23 | 12 | 24 | 20 |
| # of Control lines assayed | 12 | 12 | 12 | 12 |
| % of transfers delayed due<br>to males | 0.14 | 2.37 | 0.41 | 1.68 |
| % of transfers delayed due<br>to dormant eggs | 0.23 | 2.66 | 1.36 | 3.37 |

At the start of a size-at-birth assay, five newborn daughters of the current focal individual were taken from each MA line. A few lines did not have sufficient daughters and were excluded from the assay. This froze mutation accumulation at the current generation. At the same time, five daughters were collected from each of the exephippial individuals used for control lines. Each individual was placed in a 150-ml beaker with 100 ml of filtered lakewater and fed 20,000 cells/ml of *A. falcatus* every day; these became the grandmothers of measured individuals. We kept grandmothers and mothers under uniform conditions because maternal effects are known to be mediated through offspring size (Agrelius & Dudycha 2025). Animals were transferred to new beakers and lakewater thrice weekly (M, W, F) and were monitored for reproductive events. Ten daughters per line (2 per grandmother) were taken from the third clutch of offspring, and each placed in their own 150-ml beaker with 100 ml of filtered lakewater. Transfers and feeding for the maternal generation were the same as in the grandmaternal generation. When mothers released their second clutch, we began monitoring developmental progress of their third clutches so that the *Daphnia* mothers could be placed in filtered lakewater with no algae present immediately prior to releasing the third clutch. This ensured we were able to collect third-clutch neonates while preventing them from gathering any algae via filter-feeding and thus increasing their mass. This is a long-established procedure to measure size at birth in *Daphnia* (Tessier & Consolatti 1991; Dudycha & Tessier 1999).

For each line, we pooled third-clutch neonates from all mothers into a single beaker, then haphazardly isolated 18 neonates from the pool for measurements. Neonates were placed onto microscope slides and water was drawn off with a folded lint-free tissue to fix neonates in a position such that they would not break when removed for weighing. Neonates were dried by placing slides into a drying oven at 60°C for 48 hr and then stored in airtight containers with indicator desiccant to remove any moisture present.

Neonate dry mass was then obtained with a Mettler-Toledo UMX2 Ultra-Microbalance. We used an insect pin with an electrostatic charge (generated by brushing the pin along the investigator’s arm or eyebrow) to carefully transfer neonates from slides to the ultrabalance. Being that neonates are extremely small, we weighed the 18 neonates two-at-a-time and divided that mass by two, producing nine independent observations of neonate mass per line.

### Data analysis

We assessed normality of our data in QQ plots separated by ancestor and treatment (i.e., control vs. MA); these were reasonably linear and allowed us to proceed with parametric statistics. We used mixed-model nested ANOVA in Systat 13 to test whether our ancestors differed from each other, with control lines as the unit of replication, REML for estimation, and type III sums of squares. In this analysis, mass was the dependent variable, ancestral clone was a fixed factor, and line nested within ancestor was a random factor. Ancestral clone and line are categorical variables. We included the intercept as a fixed covariate.

We further used mixed-model nested ANOVA in Systat 13 to test whether mutation altered size at birth in each of the four independent experiments. We again had lines as the unit of replication, REML for estimation, and type III sums of squares. In this analysis, mass was the dependent variable, treatment (MA vs. ancestral control) was a fixed factor and line nested within treatment was a random factor. Treatment and line are categorical variables. We included the intercept as a fixed covariate.

To estimate the mutational variance and coefficients of variation, we calculated the variance among MA lines’ mean values. We tested for increased variance among lines following mutation accumulation in Systat 13 with an *F*-test for the equality of two variances (Snedecor & Cochran 1983).

To test whether offspring birth mass differed between individual MA lines and their respective controls, we conducted one-way ANOVAs using the lme4 package in R version 4.0.3. We applied Bonferroni corrections for multiple comparisons within each population (*n* = 12-24 tests per population).

## Results

Size at birth was not significantly different among the controls of the four ancestral clones (*F* = 1.687; *df* = 3, 44; *p* = 0.184). The estimated size of TRO was 2.83 μg, LIS was 2.61 μg, MORG was 2.48 μg, and SED was 2.45 μg (Figure 1; Table 2). These are typical masses at birth for *D. pulex* (Tessier & Consolatti 1991; Tessier, et al 1992; Dudycha & Tessier 1999; Dudycha & Lynch 2005).

**Figure 1.**
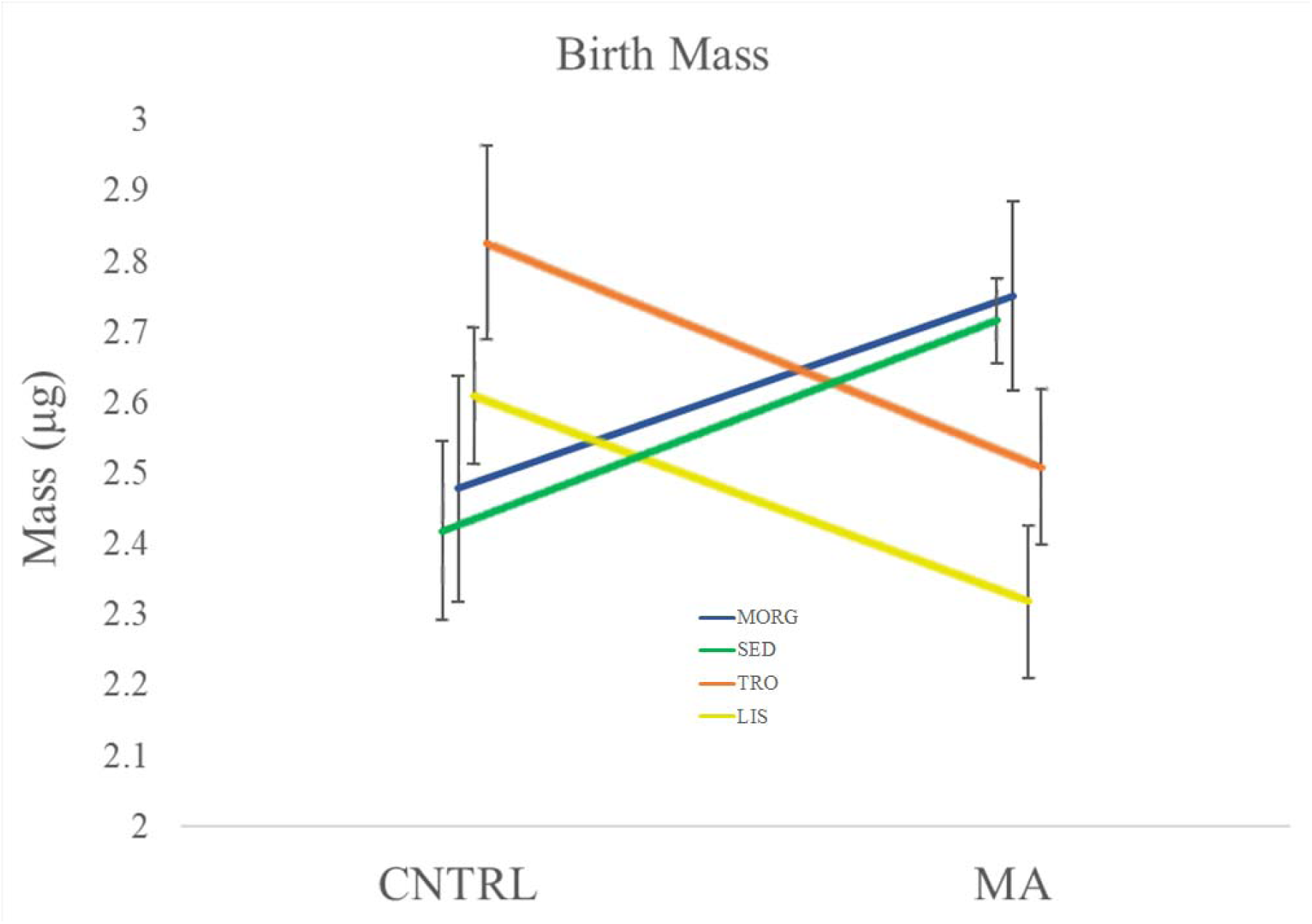
Mass at birth of *Daphnia pulex* before (CNTRL) and after mutation accumulation (MA). Error bars indicate standard error across replicate lines. Alt text: Graph comparing mass at birth of ancestral controls and descendant mutation accumulation lines for four independent mutation accumulation experiments, showing mean and standard error.

**Table 2.** Analysis of the birth mass for the four clones of *D. pulex.* All values are estimated by treating *line* as the unit of replication.

|  | MORG | SED | TRO | LIS |
| --- | --- | --- | --- | --- |
| **Control Lines** |  |  |  |  |
| Mean ($\mu\text{g}$) | 2.478 | 2.451 | 2.826 | 2.610 |
| Variance ($\mu\text{g}^2$) | 0.3071 | 0.1904 | 0.2257 | 0.1122 |
| CV | 0.2236 | 0.1780 | 0.1681 | 0.1284 |
| **MA Lines** (Avg Generation) | (93.43) | (72.25) | (86.54) | (52.20) |
| Mean ($\mu\text{g}$) | 2.751 | 2.749 | 2.509 | 2.319 |
| Percent Change | 11.02% | 12.16% | -11.22% | -11.15% |
| Variance ($\mu\text{g}^2$) | 0.4087 | 0.0436 | 0.2895 | 0.2343 |
| CV | 0.2324 | 0.0760 | 0.2145 | 0.2088 |
| **Per Generational Change** |  |  |  |  |
| Mean ($\Delta\text{M}$) | 0.0029 | 0.0041 | -0.0037 | -0.0056 |
| Variance ($\Delta\text{V}$) | 0.0011 | -0.0020 | 0.0007 | 0.0023 |
| **$\Delta\text{M}$ Tests** (MA v Control) |  |  |  |  |
| *p*-value | 0.219 | **0.0441** | 0.0923 | 0.0773 |
| *F*-statistic | 1.570 | 4.558 | 3.001 | 3.347 |
| *df* | 1, 33 | 1, 22 | 1, 34 | 1, 30 |
| **$\Delta\text{V}$ Tests** (MA v Control) |  |  |  |  |
| *p*-value | 0.638 | **0.022** | 0.686 | 0.212 |
| *F*-statistic | 0.751 | 4.364 | 0.780 | 0.479 |
| *df* | 11, 22 | 11, 11 | 11, 23 | 11, 19 |

For TRO and LIS, mass estimates for neonates of MA lines were on average smaller than those of the control lines, but this was only marginally significant (p < 0.1) in either TRO or LIS (Figure 1; Table 2). The reverse was true for lines of MORG and SED, and here only SED produced a statistically significant result (Figure 1; Table 2). Regardless of the direction of change or average number of generations, mutation accumulation altered estimated mean neonate size by 11-12% (Table 2). This equates to changing neonate size by 0.003-0.005 μg per generation (Table 2).

Estimated variance among the MA lines was greater than among the control lines for TRO, LIS, and MORG, but these estimated differences were not statistically significant (Table 2, Figure 2). Contrary to expectations, variance in the SED MA lines was significantly lower than in the corresponding control lines; in fact, the variance of the SED MA lines was an order of magnitude lower than any of the other MA lines (Table 2). The high number of line extinctions in SED may explain both the low variance and the statistical significance, since they coincidentally led to a balanced statistical design in the SED lines. Despite the lack of evidence for a general increase of variance, in TRO, LIS, and MORG lines, some individual MA lines were larger than the ancestors, while others were smaller (Figure 2, Table 3). In no case was there a balanced distribution of larger and smaller MA lines.

**Figure 2.**
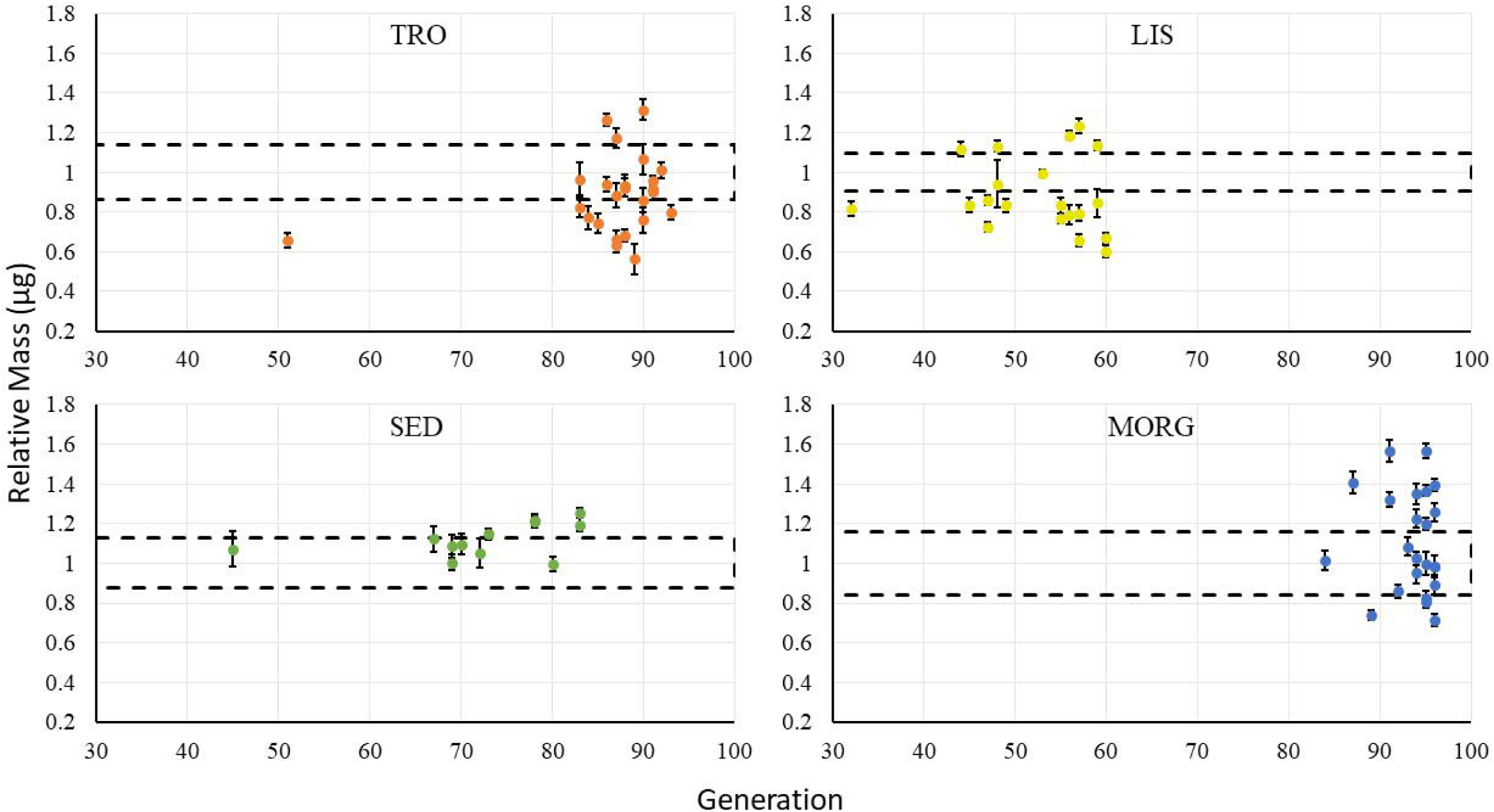
Relative mass of MA lines compared to Control lines. In each panel, mean control mass is scaled to 1; dashed lines show the standard error of the control lines’ mean. Error bars on points show standard error for each individual MA line. Alt text: Four graphs illustrating the mass at birth and number of generations of mutation accumulation for each individual line, with standard errors. Mass is given relative to the mass at birth of the ancestral control. Dashed lines across the graphs illustrate the range of control lines standard error of the mass at birth.

**Table 3.** Statistical tests of whether each individual MA line is different from the ancestral control lines’ mean. Bolded *p*-values are significant at the 0.05 level or better after Bonferroni correction. Shading indicates the direction of difference: MA lines with red shading have smaller mass at birth than the ancestor, blue shading indicates larger size at birth in the MA line. Pale shading indicates marginal statistical significance (*p* < 0.1).

| TRO |  |  | MORG |  |  | LIS |  |  | SED |  |  |
| --- | --- | --- | --- | --- | --- | --- | --- | --- | --- | --- | --- |
| Line | F-statistic | p-Value | Line | F-statistic | p-Value | Line | F-statistic | p-Value | Line | F-statistic | p-Value |
| T1 | 30.0821 | 0.000654 | M1 | 0.040012 | 1 | L2 | 8.074521 | 0.208657 | S1 | 3.356695 | 0.991861 |
| T2 | 8.401565 | 0.220994 | M2 | 6.757579 | 0.404862 | L3 | 76.2248 | 0.000000 | S2 | 0.938989 | 1 |
| T3 | 34.4981 | 0.000282 | M3 | 17.18989 | 0.012624 | L4 | 54.11668 | 0.000011 | S7 | 12.8653 | 0.023598 |
| T4 | 1.619467 | 1 | M4 | 1.873471 | 1 | L5 | 57.87057 | 0.000006 | S9 | 9.832546 | 0.065326 |
| T5 | 20.60496 | 0.005383 | M5 | 0.100907 | 1 | L6 | 9.185774 | 0.137517 | S10 | 0.004345 | 1 |
| T7 | 2.324689 | 1 | M6 | 23.07423 | 0.00284 | L7 | 12.48596 | 0.044393 | S11 | 0.550788 | 1 |
| T8 | 4.502192 | 1 | M7 | 0.320286 | 1 | L9 | 8.742306 | 0.162011 | S12 | 1.674428 | 1 |
| T9 | 18.4306 | 0.00942 | M8 | 0.040118 | 1 | L10 | 25.99625 | 0.001276 | S16 | 17.53456 | 0.005993 |
| T10 | 0.999226 | 1 | M9 | 19.39496 | 0.007014 | L12 | 19.21383 | 0.006392 | S17 | 5.767227 | 0.320655 |
| T11 | 2.866645 | 1 | M10 | 10.81398 | 0.088883 | L13 | 0.697044 | 1 | S18 | 2.255619 | 1 |
| T12 | 0.030565 | 1 | M11 | 53.99375 | 0.000013 | L14 | 6.329091 | 0.420578 | S22 | 0.00178 | 1 |
| T13 | 0.604435 | 1 | M14 | 8.36484 | 0.214757 | L17 | 17.70401 | 0.009539 | S25 | 11.7649 | 0.033687 |
| T15 | 11.8823 | 0.064819 | M15 | 6.506375 | 0.449134 | L18 | 9.716671 | 0.113476 |  |  |  |
| T16 | 3.415837 | 1 | M16 | 3.372696 | 1 | L19 | 0.017588 | 1 |  |  |  |
| T17 | 1.190196 | 1 | M17 | 1.172857 | 1 | L20 | 12.65936 | 0.041994 |  |  |  |
| T18 | 8.292502 | 0.230343 | M18 | 0.038911 | 1 | L21 | 26.88232 | 0.001054 |  |  |  |
| T20 | 0.990474 | 1 | M19 | 11.93685 | 0.061015 | L22 | 19.53474 | 0.005883 |  |  |  |
| T21 | 0.275579 | 1 | M20 | 0.000916 | 1 | L23 | 13.50278 | 0.032205 |  |  |  |
| T22 | 26.96261 | 0.001243 | M22 | 25.99102 | 0.001469 | L24 | 15.45816 | 0.017912 |  |  |  |
| T23 | 38.76012 | 0.000134 | M23 | 26.66491 | 0.00127 | L25 | 37.6793 | 0.000134 |  |  |  |
| T24 | 32.27681 | 0.000427 | M24 | 5.194071 | 0.7911 |  |  |  |  |  |  |
| T25 | 14.47958 | 0.028692 | M26 | 13.8039 | 0.033751 |  |  |  |  |  |  |
| T29 | 44.60628 | 0.000053 | M29 | 50.94411 | 0.000020 |  |  |  |  |  |  |
| T30 | 5.093849 | 0.863535 |  |  |  |  |  |  |  |  |  |

## Discussion

Size at birth mutated in different directions in different genetic backgrounds in our experiment, although we also found considerable stochasticity of mutation direction within a single genetic background. Overall, statistical support for directional bias within a genetic background was modest. Examining individual lines, however showed that two sets (TRO and LIS) had substantially more MA lines that produced offspring smaller than the ancestor, while the other two sets (SED and MORG) had more lines that produced large offspring. Overall, out of our 79 total mutation accumulation lines reflecting a net 6136 generations of mutation, 19 lines experienced significant decreases of size at birth and 14 lines experienced significant increases. Unlike fitness itself (Lynch & Walsh 1998 (Ch. 12), Bank, et al 2014), complex traits whose fitness consequences are determined by ecologically mediated tradeoffs are not necessarily expected to show overall directional effects of spontaneous mutation. Given that directionality trends differed among ancestors, and that other similar species can have either larger or smaller neonates (e.g., Ebert 1991; Dudycha & Lynch 2005), we think it is unlikely that intrinsic physiological properties constrained mutation to a particular direction. To the extent that we saw differences, different genetic backgrounds may simply have different mutational space available to them at the scale of 50-100 generations. We did not observe any association between the number of generations a set of lines had mutated prior to being assayed and the outcome of the assays (Figure 2) but it is unlikely that the timescale we used exhausted the full range of potential mutational space.

Under standard laboratory conditions, *Daphnia* are freed from nearly all selective constraints: they have no predators or pathogens, food is abundant, temperatures are benign, and ultraviolet radiation is not an issue. In this environment, offspring should be able to survive above some physiologically minimal investment by a mother, and her fitness would be maximized by producing large clutches of tiny offspring. Stressful environments often induce mothers to package their reproductive output in smaller clutches of larger offspring (Ocampo, et al. 2012; Agrelius & Dudycha, 2025). Some authors have argued that spontaneous mutations can be thought of as a type of stress upon organismal physiology (Agrawal & Whitlock 2010), eliciting phenotypes similar to plasticity induced by environmental stress. This is an unsatisfactory explanation for our results. Even though two of our sets had many lines that mutated toward larger offspring size— which is the phenotypic shift expected for most types of stress in *Daphnia* (reviewed in Agrelius & Dudycha 2025)—two sets primarily showed lines mutate in the opposite direction.

Broad- and narrow-sense heritabilities for size-at-birth in *Daphnia* are typical for life history traits. Pfrender & Lynch (2000) reported a narrow-sense heritability of 0.25 and broad-sense heritabilities of length-at-birth ranging from 0.18 to 0.37 in *D. pulex.* In an experiment with a similar operational protocol to what we report here, Dudycha & Tessier (1999) reported more than two-fold variation of mass-at-birth among clones of *D. pulex*. Ebert, et al (1993) estimated broad-sense heritabilities of length-at-birth in *Daphnia magna* to range from 0.3 to 0.6 (depending on clutch) under high-food conditions, and 0.2 to 0.4 under low-food conditions. Their estimates of narrow-sense heritabilities were similar, although somewhat higher under low-food conditions. Other species that have been shown to harbour genetic variation of offspring size include *D. pulicaria* and *D. parvula* (Tessier & Consolatti 1989). Although we do not know which genes actually contribute to the determination of offspring size, there appears to be ample genetic control over size-at-birth in *Daphnia* to expect mutation to alter offspring size.

Phenotypic plasticity also contributes to variation of offspring size in *Daphnia*, and can be influenced by maternal ontogeny, crowding, food levels, temperature, and predator exposure (Tessier & Consolatti 1989; Lüning 1992; Glazier 1992; Burns 1995; Reinikainen and Repka 2003; Guinnee, et al. 2007; Machácek and Seda 2013; Gabsi, et al. 2014). Many researchers studying phenotypes in *Daphnia* avoid using individuals from the first clutch (and sometimes second, as we do) as experimental animals due to visibly different sizes of neonates from the initial clutch (Glazier 1992). It is for these reasons that we maintained mothers and grandmothers of experimental animals under standardized conditions of density, temperature, food level, and predator exposure, attempting to minimize maternally driven phenotypic plasticity.

We were surprised to see that our mutation accumulation lines did not have greater among-line variances than they did, given that by definition mutation creates new variants, though individual lines clearly differed from the ancestors. Most mutation accumulation experiments show an increase of variance for complex traits (e.g., Lynch, et al 1999; Baer, et al. 2005, 2007; Latta, et al 2013, 2015; Davenport, et al. 2021). Several possibilities could explain why we did not find this. First, our replicate control lines were not the ancestor itself, and ∼6 generations separated the assayed individuals from the ancestor. However, a dramatic increase of mutational variation in early generations followed by no further increase in the much larger number of generations that came later is implausible. Given that three of our ancestors’ mutational descendents were trending towards greater among-line variances, it is also possible that greater replication (or a balanced design) may have yielded stronger statistical confidence in those increases.

Some might suggest that our experiment may have provided an inadequate opportunity for mutation to occur and produce new genetic variation of mass at birth. However, a paucity of total mutation is unlikely to be an explanation for the patterns we saw. First, an earlier report of life history effects in one set of the lines much earlier than reported here (the LIS lines at generation 37; Davenport, et al 2021) found clear mutational variance. Second, Keith, et al (2016) estimated the base substitution rate of *Daphnia pulex* to be 5.7 x 10 per site per generation. Ye, et al (2017) estimated the genome size of *D. pulex* clone PA42 to be ∼156,400,000 bp in comparison to clone TCO at ∼197,300,000 bp. Given the average number of generations and numbers of lines, this implies that our four experiments contained somewhere between 773 (the low estimate for SED) and 2416 base substitutions (the high estiamte for MORG). However, these figures are underestimates of the potential fuel for change to quantitative traits, because they exclude other types of mutations that may have bigger impacts on phenotypes. Keith, et al (2016) also observed many large-scale duplications and deletions, yielding gene duplication and deletion rates of 3.27 x 10 and 3.71 x 10 per gene. Given estimates of gene numbers in *D. pulex* that range from 18,440 to 30,097 (Ye, et al 2017), that implies 523 to 2114 gene duplications and 594 to 2399 gene losses in our experiments. All told, this means that 6.06 – 14.99% of all protein-coding loci were hit by either a duplication or a loss in a given experiment. Further fuel would come from small indels affecting protein-coding loci or regulatory regions, though the potential impact of these is hard to assess in the absence of information on the size of regulatory regions.

Finally, an interesting real possibility has been raised by Johnson, et al. (2020). This is that mutation accumulation experiments may reveal epigenetic variation when they assay replicated control lines. Since the genetic variation among such lines should be zero, what should we make of the observed among-line variance that did occur? One might turn to ordinary stochasticity as an explanation, but given that we had substantial within-line replication, this seems unlikely. However, epigenetic variation is possible, potentially tied to each unique individual we hatched out of control ephippia. This would imply that there is substantial epigenetic variation for size at birth in *D. pulex* and it is not overtaken by mutational variation in the course of 50-100 generations.

In some work, researchers have estimated mutational variance in experiments that lack assays of replicate control lines (reviewed in Conradsen, et al. 2022). In these cases, the quite reasonable assumption is that there simply was no ancestral genetic variation, and that whatever among-line variance is detected in a set of MA lines has been an increase from zero. Had we approached our data that way, and compared the MA lines against just the mean ancestral value, we would have inferred per-generation increases in mutational variance that were roughly quadruple what we observed, and seen statistically significant directional bias in each set of MA lines. Given our results, it is worth investigating Johnson’s et al. (2020) hypothesis more directly.

## Acknowledgements

David Cann and Jake Swanson provided assistance propagating the MA lines. Ancestral clones MORG5, LIS3, and TRO3 were originally provided by Mike Lynch. Brian Hollis, Carrie Wessinger, Arnaud Le Rouzic, and two anonymous reviewers provided thoughtful comments on earlier versions that allowed us to improve this work. Funding for this work was provided by grant DEB-1556645 from the U.S. National Science Foundation to JLD.

